# Pregnancy and weaning regulate human maternal liver size and function

**DOI:** 10.1101/2021.02.18.431862

**Authors:** Alexandra Q Bartlett, Kimberly K Vesco, Jonathan Q Purnell, Melanie Francisco, Erica Goddard, Andrea DeBarber, Michael C Leo, Eric Baetscher, William Rooney, Willscott Naugler, Alex Guimaraes, Patrick Catalano, Pepper Schedin

## Abstract

**BACKGROUND:** During pregnancy, the rodent liver undergoes hepatocyte proliferation and increases in size, followed by weaning-induced involution via hepatocyte cell death and stromal remodeling, creating a pro-metastatic niche. These data suggest a mechanism for increased liver metastasis in postpartum breast cancer patients.

**OBJECTIVES:** Investigate if the human liver changes in size and function during pregnancy and weaning.

**METHODS:** Abdominal imaging was obtained in healthy women at early and late pregnancy, and post-wean. During pregnancy time points, endogenous glucose production was measured and fasting blood taken to measure bile acids.

**RESULTS:** Independent of weight gain, most women’s livers increased in size with pregnancy, returning to baseline post-wean. Putative roles for bile acids in liver growth were observed in pregnant women and rodents.

**CONCLUSIONS:** The human liver is regulated by reproductive state with growth during pregnancy and volume loss post-wean. These findings may have broad implications for sex-specific liver diseases and cancer.

## Introduction

Sex-specific differences in liver disease have been attributed to sexual dimorphisms in steroid production, metabolic enzymes, and behavior patterns *(1)*. Whether a pregnancy cycle contributes to sex-specific liver disease remains largely unexplored. A previously unrecognized liver biology linked to reproductive status has been reported in rodents. During pregnancy and lactation, hepatocytes proliferate and enter into a higher anabolic state accompanied by an overall increase in liver size. Upon weaning, hepatocytes undergo programmed cell death, liver metabolism shifts towards catabolism, and the liver regresses to its pre-pregnant size in a process referred to as weaning-induced liver involution *(2)*. In mice, liver involution promoted breast cancer metastasis to the liver, suggesting a pathophysiological consequence of liver involution *(2)*. Notably, young women diagnosed with breast cancer in the postpartum period were found to be at increased risk for liver metastasis *(2)*. Taken together, these findings suggest that pregnancy, lactation, and weaning may create a pro-metastatic niche in the human liver *(2)*. However, it is unknown whether the human liver changes in size and function across a reproductive cycle, as expected if the liver is tuned to meet the unique metabolic demands of reproduction. Such evidence would corroborate the findings in rodents and be foundationally important for future studies of liver health in women.

To investigate the impact of reproductive state on liver size and function in women, we conducted a prospective study of healthy pregnant women using magnetic resonance and spectroscopy imaging of the liver and compared findings to a validated rodent model. Here, we show that the human female liver is regulated in both size and function by reproductive state, and provide the first ever evidence of weaning-induced liver involution. Further, these data provide a novel hypothesis to explain the increased liver metastasis observed in postpartum breast cancer patients, as well as having potentially broader implications for the understanding of sex-specific liver diseases.

## Results

Forty seven healthy pregnant women completed early (12-16 weeks gestation) and late (32-36 weeks gestation) pregnancy study visits **(Fig 1A)**, where they underwent liver magnetic resonance imaging (MRI) (**Fig A’**), provided blood samples, and completed body composition analyses. Participant demographics are shown in **STable 1**. On average, liver volumes increased 15% (182 cm^3^ ± 197 cm^3^) from early to late pregnancy *(P* <0.001) (**Fig 1B**). Because liver size is attuned to overall body size via the “hepatostat” *(3)*, we determined whether increase in liver size from early to late pregnancy correlated with increased body mass. First, we investigated the existence of the hepatostat at baseline, using body weight at the early pregnancy visit as a surrogate, as pregnancy-related weight gain is minimal at this time point *(4)*. Liver volumes at early pregnancy correlated with body weight (**Fig 1C**), confirming previous studies in non-pregnant individuals *(3)*. In contrast, the change in liver volume during pregnancy did not correlate with gestational weight gain (**Fig 1D**). We also found no relationships between pregnancy liver volume change and change in total fat mass (**Fig 1E**), subcutaneous abdominal- or visceral-adipose tissue volumes (**Table 1**). However, we found that the change in a woman’s *fat free* mass, which includes liver, fetal tissue, placenta, and plasma correlated with change in liver size (**Fig 1F)**. These data suggest that liver size increase during pregnancy is unlinked to overall body size (i.e. not by hepatostat regulation), but rather may reflect an unrecognized reproductive state-controlled program.

**Table 1.**
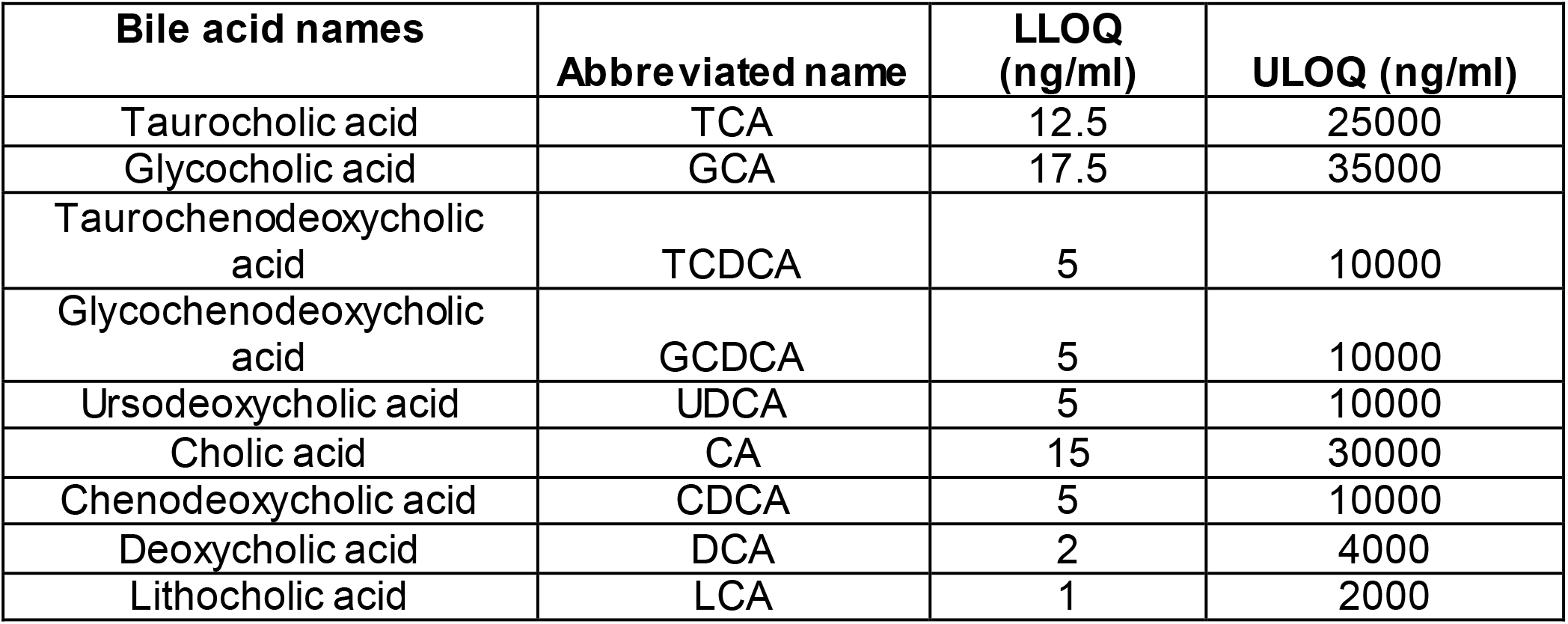
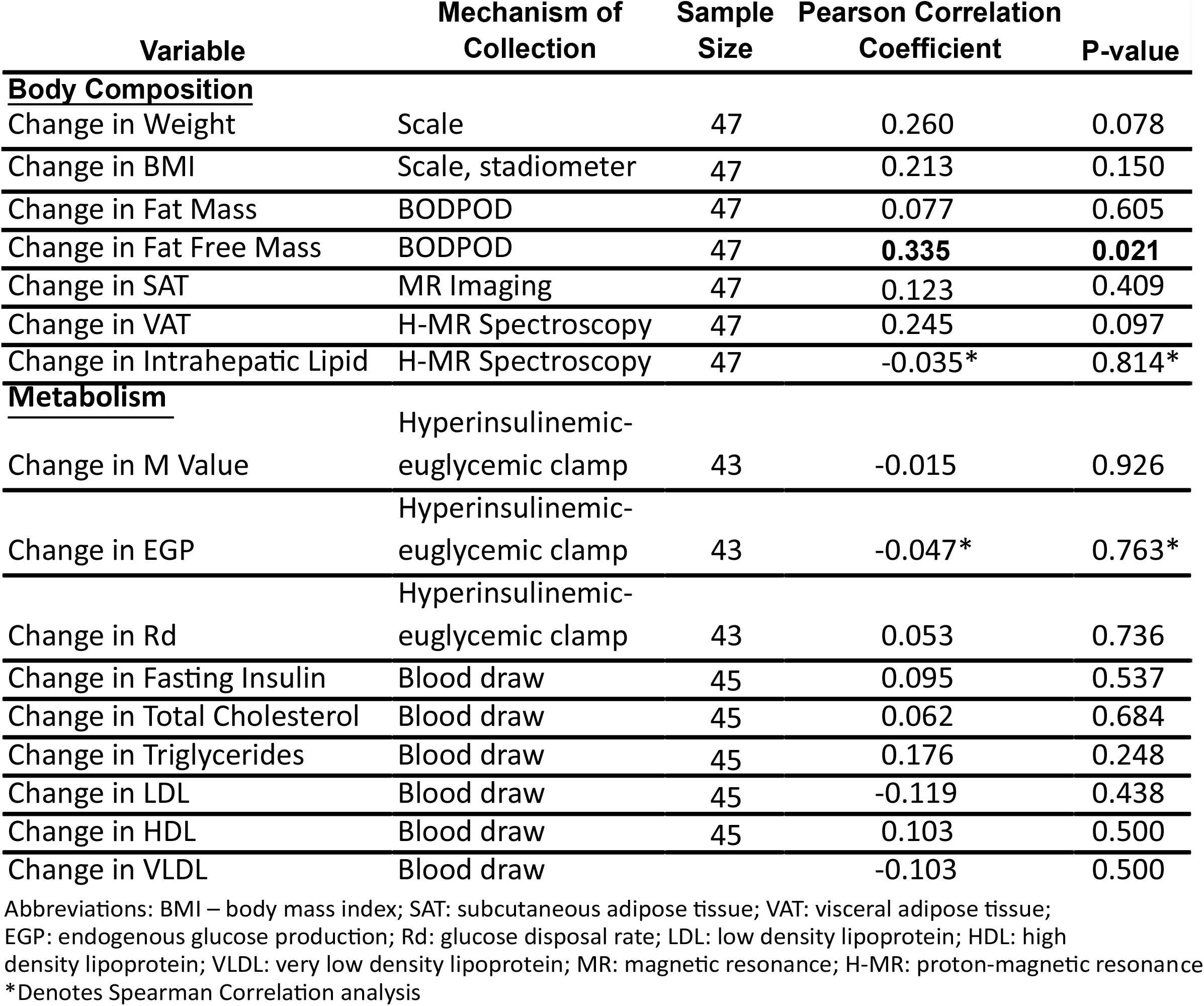
Change in liver volume correlated with measures of body composition and metabolism.

**Figure 1.**
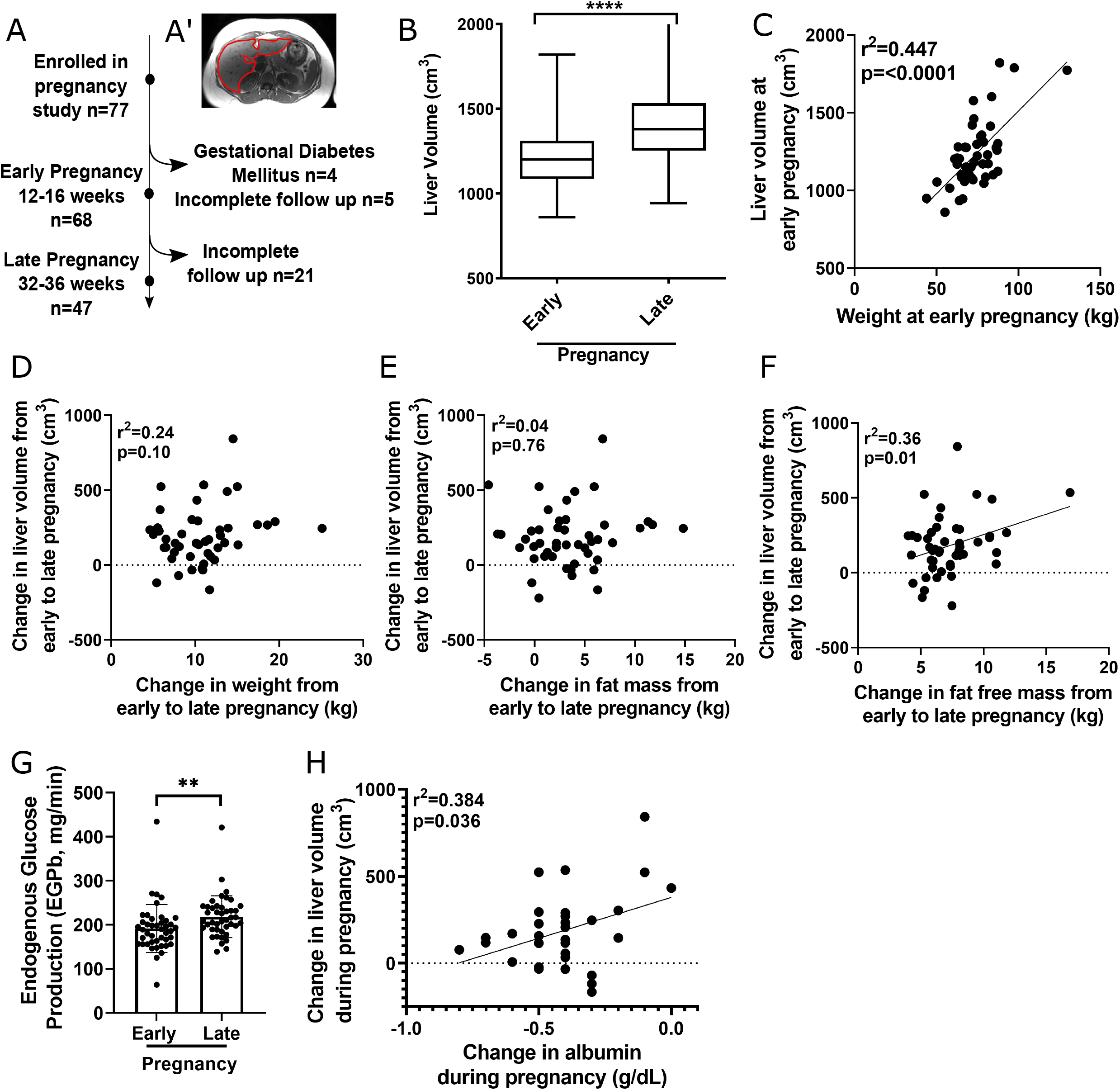
Liver changes during pregnancy: (A) Diagram of the observational study. (A’) Liver MRI cross-section. (B) Average liver volume at early and late pregnancy (P: **** < 0.0001 by Two-tailed paired T test). (C) Pearson’s correlation of liver volume and BMI at early pregnancy (N = 47). Pearson’s correlation of change in liver volume with change in body weight (D), fat mass (E), and fat free mass (F). (G) Endogenous glucose production (EGP-b) at early and late pregnancy (P: ** < 0.01 by Two-tailed paired T test). (H) Pearson’s correlation of change in liver volume and change in albumin (N = 30).

We next asked if metabolic measures were associated with liver volume change and found no relationship with cholesterol concentrations or with measures of insulin sensitivity, i.e. endogenous glucose production (EGP) and glucose disposal rate (Rd) (**Table 1**). We also found no relationship between change in liver volume and change in intrahepatic lipid content (**Table 1**). Assessment in rodents also showed no change in intrahepatic lipid during pregnancy (**SFig1**). In sum, we observed that the increase in human liver volume with late pregnancy occurred independent of weight gain of pregnancy, various other measures of body composition, circulating metabolites, and intrahepatic lipid storage.

Increased liver size during pregnancy could indicate increased function. We found evidence for increased liver output at late pregnancy as measured by EGP and albumin concentration, surrogates of liver function (**Fig1G and 1H**). These data are consistent with an increase in liver synthetic capacity during pregnancy.

We next looked for evidence of weaning-induced liver involution in women, a biology not previously described in humans. Of the 47 women who participated in our pregnancy study, 36% (n=17) completed a liver MRI >3 months post-wean (median 5.7 months) (**Fig 2A)**. Liver volumes trended toward a decrease in size between late pregnancy and post-wean (**Fig 2B**). Further, post-wean liver volumes were similar to early pregnancy, indicative of a return to baseline (**Fig 2C**). These data provide the first evidence suggesting liver involution in women.

**Figure 2.**
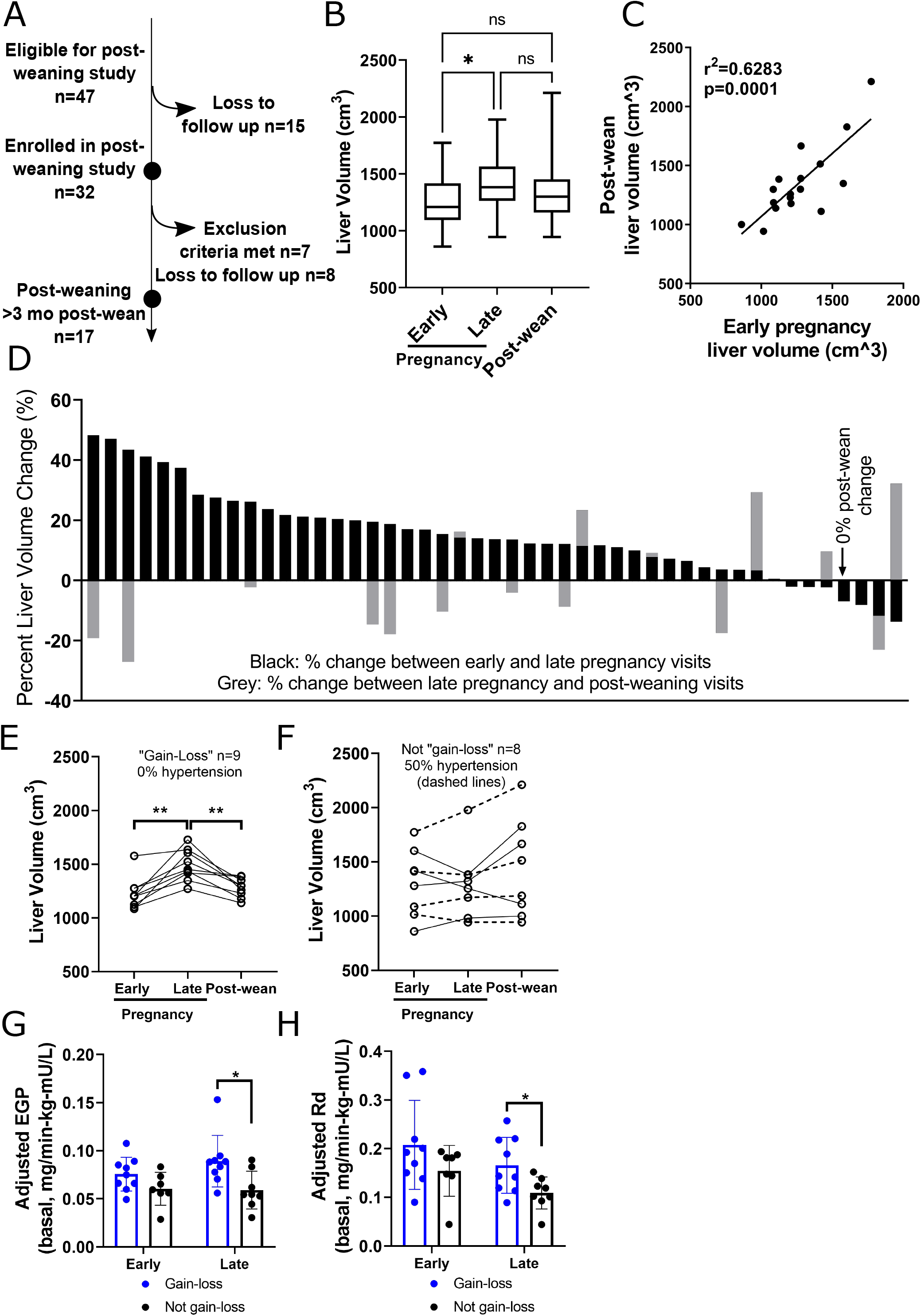
Human liver volumes post-wean: (A) Diagram for post-wean observational study. (B) Liver volume at early and late pregnancy, and post-wean time points (N=17). (C) Pearson’s correlation of liver volumes at early pregnancy and post-wean (n=17). (D) Liver volume change between early and late pregnancy (black bars) and between late pregnancy and post-wean (grey bars) per participant. Primary (E) and secondary patterns (F) of liver volume change with pregnancy and post-wean. Dashed lines show participants with hypertension (Paired T-test). Endogenous glucose production (EGP) (G) and glucose disposal rate, Rd, (H) in women in gain-loss group compared to women in not gain-loss group, paired T test. P value: * < 0.05, ** < 0.01.

While our data showed a statistically significant increase in liver size during pregnancy and a trend toward decrease after weaning, there was heterogeneity in how individuals’ liver sizes changed with pregnancy and post-wean. During pregnancy, we found that 72% (34/47) of women had a 20% average increase in liver volume (**Fig2D, black bars**). However, 21% of participants (10/47) had no measurable liver volume change and 6% (3/47) had a reduction in liver volume (**Fig 2D, black bars, STable 2)**. We saw similar heterogeneity with regard to liver volume change from late pregnancy to post-wean (**Fig 2D, grey bars**).

Considering the heterogeneity in liver volume change, as well as what is known of normal rodent liver biology (i.e. liver weight gain with pregnancy and loss post-wean) *(2)*, we performed subgroup analyses. We delineated the participants into two groups: “*gain-loss*”, the observed pattern in the normal rodent, or “*not gain-loss*” for those that did not display this expected pattern. 53% (9/17) of women displayed the anticipated liver gain-loss pattern (**Fig 2E**). The not gain-loss group (8/17) comprised heterogeneous patterns and included three women who lost liver volume during pregnancy and re-gained post-wean, three women with no significant liver volume changes, and one woman each with either continuous liver size loss or gain across the 3 visits (**Fig 2F**). Of note, the liver volume patterns of gain-loss and not gain-loss did not associate with a woman’s overall weight gain of pregnancy (**SFig 2**).

Upon further exploration, we found that none of the gain-loss group had gestational hypertension, yet 50% of the not gain-loss group did (**Fig 2F, dashed lines**). Further, measures of insulin sensitivity differed between these groups. Specifically, we found the gain-loss participants had greater EGP at late pregnancy (**Fig2G**), consistent with published data showing elevated EGP in healthy pregnancy *(5)*. We also found greater glucose disposal rate at late pregnancy in the gain-loss group (**Fig 2H**), consistent with greater insulin sensitivity in the muscle. These data suggest that the not gain-loss pattern may be associated with suboptimal gestational metabolic health and gestational hypertension. One question is whether these metabolic parameters impact fetal outcomes. In this cohort, maternal liver gain-loss patterns did not correlate with newborn weight, length, or Ponderal index, three common neonatal health measures.

To investigate the mechanistic relationship between reproductive state and liver size, we utilized tissues from a previously published rat pregnancy study *(2)*. We found liver weight increases during pregnancy were greater than expected due to gestational weight gain alone, indicating rat liver weight during pregnancy is unlinked from the hepatostat, corroborating our human data (**Fig 3A**). Next, we confirmed maximum hepatocyte proliferation during pregnancy **(Fig 3B)**, consistent with previous reports *(2, 6)*. Together, these data suggest a physiological model in which increased liver volume of pregnancy is due to increased hepatocyte proliferation that is activated via an unrecognized, pregnancy-mediated developmental program.

**Figure 3.**
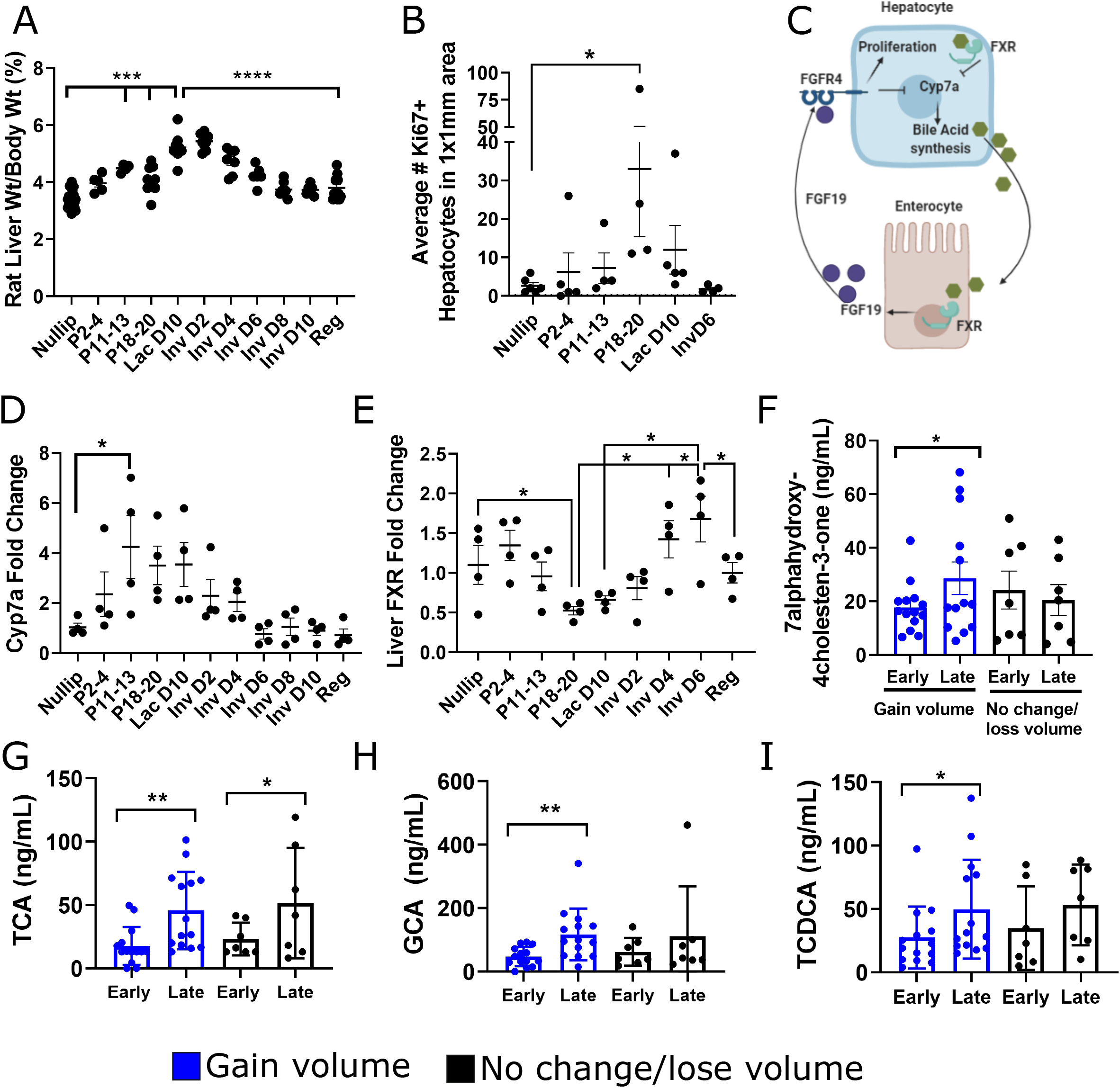
Hepatic bile acid signaling and liver size: (A) Rat liver weight normalized to body weight: Nulliparous (nullip) n=24; early (P2-4) n=5; mid (P11-13) n=4; and late (P18-20) pregnancy n=10; lactation day 10 (Lac D10) n=9; involution (Inv) day 2 n=9; InvD4 n=7; InvD6 n=6; InvD8 n=7; InvD10 n=6; Regressed (Reg) n=14; One-way ANOVA. (B) Ki67+ hepatocytes in rat livers, n=3-5/group. (C) Bile acid signaling model. Cyp7a (D) and Fxr (E) mRNA fold change in liver, n=4 per group; One-way ANOVA. (F) Human 7α-hydroxy-4-cholestene-3-one (7αC4) plasma concentrations at early and late pregnancy, separated by liver gain (n=14) and no gain patterns (n=7). Plasma concentrations of bile acids TCA (G), GCA (H), and TCDCA (I) Paired T test, P value: * < 0.05, ** <0.01, *** < 0.001, **** < 0.0001.

As a possible mechanism underlying a pregnancy-associated liver growth program, we investigated bile acid metabolism in our rat model, as bile acid signaling contributes to liver regeneration following partial hepatectomy and can control liver size independent of body size *(7–9)*. Further, bile acids have been shown to regulate hepatocyte proliferation (**Fig 3C**) *(7, 10)*. To investigate if the bile acid pool is modulated by reproductive state, we measured liver *Cyp7a*, a rate limiting enzyme in bile acid synthesis *(7)*. We found 3-4 fold increased expression of *Cyp7a* with pregnancy, with expression remaining elevated during lactation, followed by a rapid decline with weaning (**Fig 3D**). Since hepatic FXR signaling acts as a negative regulator of Cyp7a **(Fig 3C**) *(8)*, we measured hepatic *FXR* and found it was downregulated during late pregnancy, when *Cyp7a* was high, and increased with weaning, when *Cyp7a* was low (**Fig 3E**). These data associate increased bile acids with the physiologic expansion of the liver during pregnancy—consistent with a previous report *(8)*—and extend these observations to suggest a role for bile acids in regulating liver size during lactation and weaning.

We then examined associations between liver growth and the bile acid pool in pregnant women by measuring a biomarker of bile acid production and plasma bile acid concentrations at early and late pregnancy. Plasma concentrations of 7α-hydroxy-4-cholestene-3-one (7αC4), a readout for cholesterol 7α-hydroxylase (Cyp7a1) enzyme activity *(11)*, were significantly increased at late compared to early pregnancy only in the women who had increased liver volume during pregnancy (**Fig 3F**). This finding supports the hypothesis that increased bile acid production during pregnancy may be required for liver size increase. Further, among these women, we found increases in several primary bile acids and their conjugates (**Fig 3G-I**). Of note, changes in secondary bile acids, which are metabolic products of gut bacteria, only weakly correlated with liver volume change (**STable 3**). In this human cohort, we did not find associations between concentration of plasma FGF19, an enterocyte product shown to induce hepatocyte proliferation and liver growth in rodents (**SFig 3**). One potential caveat to this analysis is that plasma concentrations of FGF19 may not reflect concentration in the portal vein that links the gut and liver. In sum, these human data are consistent with an increased bile acid pool during pregnancy, which may contribute to the increased liver size observed in pregnancy.

## Discussion

We report a new human liver biology in normal women, specifically increased liver size with pregnancy and decreased size post-wean, putatively to accommodate the dramatic changes in metabolic demands across a pregnancy-lactation-wean cycle. Further, in our animal studies we found evidence for a potential mechanism, bile acid signaling, for which we found supportive evidence in our human study. We speculate that liver volume changes in women who do not follow the gain-loss pattern may indicate a suboptimal response to pregnancy, as the population without this pattern was enriched for hypertension and lacked the expected change in late pregnancy insulin sensitivity *(5)*. These data were generated in a small, predominately White, non-Hispanic cohort and require validation in a larger study with a diverse population to generalize these findings.

These novel observations demonstrating reproductive control of liver size and function in women concur with recent observations in rodents, suggesting a conserved liver biology. The question of whether this newly described liver biology has implications for maternal health during pregnancy or sex-specific risk for liver disease remains to be determined *(1, 12, 13)*. However, our evidence suggestive of weaning-induced liver involution in women may lead to improved understanding of the high rates of liver metastasis observed in young postpartum breast cancer patients *(2)*.

## Methods

### Recruitment

#### Prospective cohort

We conducted a prospective cohort study of pregnant women receiving care at Kaiser Permanente Northwest (KPNW), or Oregon Health & Science University (OHSU). Recruitment started in December 2014 and was completed in August 2017. KPNW members who met the study inclusion criteria were identified weekly using the electronic health record (EHR). Eligible participants were mailed a recruitment letter and received a follow-up phone call a week later. During this telephone call, study personnel conducted additional eligibility screening and scheduled an explanatory visit. If patients consented at the explanatory visit, this was followed by two visits between 12-16 weeks of gestation and two visits between 32-26 weeks of gestation

#### Inclusion/Exclusion criteria

Patients were eligible for the study if they were between 18-45 years of age; were less than 14 weeks pregnant with a singleton gestation at time of enrollment; had a BMI between 18.5 kg/m2 and 38 kg/m^2^; and were fluent English speakers. Participants were excluded if they had any of the following conditions or symptoms: contraindications to MRI study (e.g., claustrophobia, metal implants); pregestational diabetes; gestational diabetes; history of bariatric surgery or other medical conditions requiring specialized nutritional care; anemia; current history of drug, tobacco, or alcohol use; maternal rheumatologic or chronic inflammatory state; or chronic hypertension.

### Measures

Data for this paper were collected at two study visits: one at 12-17 weeks of gestation, and one at 32-36 weeks of gestation. Height was measured at the first visit to allow for calculation of BMI; weight was measured using a calibrated scale at each visit. Demographic variables, including parity and preconception BMI, were extracted from the electronic medical record.

### Liver Volume Determination

Magnetic resonance imaging was performed on a Siemens 3T Vereo using an 8 channel phased array torso body coil. The protocol included a volume interpolated 3D gradient echo sequence for 3D volume estimates of the liver (256×192 encoding matrix, asymmetric field of view 30 x 24, TE/TR 2.4/5ms, and voxel size 1.17 x 1.17 x 2.5mm). Image analysis was performed using OsiriX (OsiriX Imaging Software, Geneva, Switzerland) software and Image J software (National Institutes of Health). For volume estimation, 3D-VIBE (a T1 weighted FLASH technique with fat selective prepulse) sequences were used. The liver was identified on each image, and the outline of the liver tissue annotated by freehand region of interest estimation by operators trained by a body radiologist with over 10 years of experience in MRI of the liver. This allowed for the generation of a liver area on each slice. Liver volume was calculated by multiplying the estimated area of each slice by the interval between slices, summing all volumes containing liver for the total liver volume *(14)*. Liver volume determinations were performed by 2 blinded operators. Operators independently measured liver volumes for 5 cases with 2 MRI scans per case (early and late pregnancy). The observed inter-operator variability (S Table 2) was used to benchmark values that are within the range of measurement error, in this case +7 to −7 percent.

### Air Displacement Plethysmography (ADP)

Air displacement plethysmography (BOD POD, COSMED USA, Inc., Concord, CA) was used to determine participants’ fat mass, fat free mass, and percent body fat at each visit. Participants first changed into a bathing suit or spandex clothing and a swimming cap. They then sat inside the BOD POD while the air displaced by the body was measured. Results included total mass and body density. Fat mass and fat-free mass were estimated using van Raaj’s pregnancy equations to account for changes in the density of fat-free mass during pregnancy *(15, 16)*.

### Magnetic resonance imaging acquisition

Magnetic resonance imaging and spectroscopy data were collected using a Siemens Prisma Fit 3T whole-body system (Siemens Healthineers, Erlangen, Germany) at the Advanced Imaging Research Center at Oregon Health & Science University. Abdominal MR data were acquired in two stations, the first centered at umbilicus and the second centered over the sternal notch, to acquire MRI and liver MRS. Siemens flexible 18-channel array and spine array receiver coils with body-coil transmission were used. The abdominal MRI protocol included a T1-weighted gradient-echo sequence (TE = 2.5ms, TR = 140ms, flip-angle = 90 degrees, (1.25mm)^2^ in-plane resolution, 30-slices with 6mm thickness) acquired in 2-breath holds of ∼18 sec each. The liver T_1_-weighted MRI protocol was acquired with identical parameters to the abdominal T_1_ volume, but with a variable number of slices to cover the entire extent of the liver.

### Magnetic resonance imaging processing

The T_1_-weighted MRI data sets of abdomen and liver were manually spliced together with affine-transformations and overlapping slice elimination. The top of the liver and the L-4/5 intervertebral disk were identified as the upper and lower bounds, respectively, for the segmentation analysis for abdominal visceral and subcutaneous fat volumes.

Abdominal T_1_-w MRI volumes were segmented into five classes: unlabeled, subcutaneous adipose tissue (SAT), visceral adipose tissue (VAT), muscle, and organ (including all other abdominal volume). A custom Python pipeline was used to create an initial automated segmentation using inputs from the umbilicus T_1_-weighted volumes, the liver T_1_-weighted volumes, and an 11-slice manual segmentation label map, the merged T_1_-weighted MRI data set, and the affine transforms that map individual volume acquisitions to the merged image space. Manually generated uterus/placenta and liver masks were created as these two regions have high rates of false-positives for classification as adipose tissue.

Processing within the pipeline made use of the following Python libraries: Nipype *(17)*, the Advanced Normalization Tools (ANTs) *(18)*, the Insight Toolkit (ITK)*(19)*, Scikit-image *(20)*,

Scikit-learn *(21)*, and SciPy *(22)*. Following N4 bias field correction, steps in the segmentation pipeline relied upon intensity thresholding and morphological operations. The muscle mask was generated with a compact watershed algorithm seeded with the muscle mask from the 11-slice segmentation. Subcutaneous adipose tissue masking made use of the geodesic active contours (GAC) algorithm *(23)*, coupled with dilation and erosion steps to distinguish the SAT from internal VAT. Visceral adipose tissue was taken as the difference between the total adipose mask and the SAT mask. Segmentation masks output from the automated pipeline subsequently underwent slice-by-slice manual review followed by manual refinement by a single analyst (JQP) using the 3D Slicer software package to ensure accuracy of VAT and SAT masks placements.

Liver segmentation was manually conducted separately using the Osirix software program by one analyst (AQB).

### Magnetic resonance spectroscopy

Intrahepatic lipid (IHL) was measured using ^1^H single-voxel magnetic resonance spectroscopy (MRS), following MR imaging. Liver MRS voxels were positioned within the right lobe with voxel sizes ranging from 18 to 24 cm^3^.

Liver spectra were collected using a PRESS single voxel spectroscopy (SVS) sequence (TR = 5 s, TE = 30 ms, 1,024 points, 2,000 kHz spectral width). The long repetition time ensured fully relaxed water signal (99.2%), because it serves as an internal standard for quantification. Three separately-acquired MRS series were run, each during a 10-second breath hold.

MRS analysis was conducted using the AMARES time-domain fitting module within the jMRUI software program. All spectral fits were inspected and rerun with additional constraints if fitting contained errors. IHL is expressed as a proportion of primary lipid peak to water peak areas.

### Hyperinsulinemic-euglycemic clamp

Hyperinsulinemic-euglycemic clamp with co-infusion of [6,6-^2^H_2_] glucose was used to determine whole body and skeletal muscle insulin sensitivity (Rd) and endogenous glucose production (EDP) *(24, 25)*. Subjects were advised regarding a standard diet consisting of 30% of total calories from fat sources, 15% from protein, and 55% from carbohydrates for the three days before study. Following an 11-hour overnight fast, subjects were admitted to the OHSU CTRC where a hyperinsulinemic-euglycemic clamp will be performed. At 0600 hr, an intravenous catheter was placed in one arm for infusions and in the contralateral hand for blood withdrawal and warmed to 70°C using a warming mitt for sampling of arterialized venous blood. A primed constant infusion of [6,6-^2^H2] glucose (Cambridge Isotope Laboratories, Andover, MA) was infused at 0.133 ml/min and an enrichment intended to achieve ∼1.0 mol percent excess for all subjects. The basal infusion of [6,6-^2^H_2_] glucose was continued for 2hr, and plasma samples were obtained from 90 to 120 minutes to estimate basal endogenous glucose production and fasting insulin concentration. Basal endogenous glucose production (EGP) was calculated according to the steady-state equations of Steele *(26)*. At the completion of the 2-hr infusion glucose isotope, a primed, constant infusion of regular insulin at 40 mU/m^2^/min was started. Plasma glucose was maintained at 90 mg/dl for the remaining 2 hours. During the final 30 minutes of the clamp, blood samples were obtained every 5 minutes for isotope analysis. Suppression of endogenous glucose production by insulin infusion during the 2-hr clamp was estimated using the method developed by Black *(27)*. EGP, Rd, and M-value were adjusted for insulin level (mU/L) and fat free mass (kg).

### Labs

Venipuncture was used to obtain blood samples with participants in the fasting state. The following measures were assessed and run in the Laboratory Core of the Oregon Clinical and Translation Research Institute (OCTRI): comprehensive metabolic panel, lipid panel, free fatty acids, liver function tests, glucose, and insulin. Insulin was assessed by radioimmunoassay (Mercodia AB, Uppsala Sweden and glucose by a Hexokinase based colorimetric assay (Stanbio laboratory, Boerne Tx 78006).

### Bile Acid Profiling

Bile acid profiling was performed in the OHSU Bioanalytical Shared Resource/Pharmacokinetics Core. Plasma samples from early and late pregnancy were utilized to quantify plasma bile acids and 7alpha-hydroxy-4-cholesten-3-one using liquid chromatography-tandem mass spectrometry (LC-MS/MS) performed with a 4000 QTRAP hybrid triple quadrupole-linear ion trap mass spectrometer (SCIEX) operating with electrospray ionization (ESI) in the negative mode The mass spectrometer was interfaced to a Shimadzu HPLC system consisting of SIL-20AC XR auto-sampler and LC-20AD XR LC pumps. Analyte separation was achieved using a gradient HPLC method and Luna 2.5u C18(2)-HST 50×2 mm column (Phenomenex) kept at 50°C with a Shimadzu CTO-20AC column oven.

The stable isotope dilution LC-MS/MS method to quantify plasma bile acids was previously described *(28)*. In brief plasma was spiked with internal standards and bile acids were measured following protein precipitation and extraction with methanol, centrifugation and filtration of the supernatant. Calibrants were prepared in charcoal stripped matrix (SP1070 from Golden West Biological) using authentic bile acid and conjugate standards (obtained from Toronto Research Chemicals and Cerilliant).

Data were acquired using SCIEX Analyst 1.6.2 and analyzed using SCIEX MultiQuant 3.0.3 software. Sample values were calculated from calibration curves generated from the peak area ratio of the analyte to internal standard versus analyte concentration that was fit to a linear equation with 1/x weighting. Calibration curves for bile acids and conjugates covered analytical measurement ranges as outlined in Table 1.

Plasma 7α-hydroxy-4-cholesten-3-one was determined by liquid chromatography-tandem mass spectrometry (LC-MS/MS) following protein precipitation and extraction with acetonitrile. To each 100 µl sample of EDTA plasma was added 1 ng of internal standard 7α-hydroxy-4-cholesten-3-one-d7 (prepared at 0.2 ng/µl in methanol) and 300 µl of acetonitrile. The samples was vortex mixed and centrifuged at 12,000 xg for 10 min. The supernatant was removed and filtered prior to injection for analysis with LC-MS/MS.

Calibration standards were prepared across the range 1 to 100 ng/ml in charcoal stripped plasma SP1070 (Golden West Biological) using authentic 7α-hydroxy-4-cholesten-3-one (obtained from Toronto Research Chemicals).

LC-MS/MS was performed using a 5500 Q-TRAP hybrid triple quadrupole-linear ion trap mass spectrometer (SCIEX) with electrospray ionization (ESI) in the positive mode. Shimadzu HPLC system consisting of SIL-20AC XR auto-sampler and LC-20AD XR LC pumps. The 5500 QTRAP was operated with the following settings: source voltage 4500 kV, GS1 40, GS2 30, CUR 40, TEM 650 and CAD gas high.

Compounds were quantified with multiple reaction monitoring (MRM) and transitions optimized by infusion of pure compounds and as outlined in Table 2. The bold transitions were used for quantification.

**Table 2.**

| Q1mass | Q3mass | Time msec | ID | DP | EP | CE | CXP |
| --- | --- | --- | --- | --- | --- | --- | --- |
| **401.2** | **177.1** | **80** | **7α-C4** | **131** | **10** | **31** | **20** |
| 401.2 | 97 | 80 | 7α-C4 | 126 | 10 | 37 | 12 |
| **408.3** | **177.1** | **80** | **7α-C4-d7** | **156** | **10** | **31** | **14** |
| 408.3 | 390 | 80 | 7α-C4-d7 | 156 | 10 | 31 | 14 |

Analyte separation was achieved using a Gemini 3u C6-Phenyl 110A 100×2 mm column (Phenomenex) kept at 35 °C using a Shimadzu CTO-20AC column oven. The gradient mobile phase was delivered at a flow rate of 0.4 ml/min, and consisted of two solvents, A: 0.1% formic acid in water, B: 0.1% formic acid in acetonitrile. The initial concentration of solvent B was 40% followed by a linear increase to 95% B in 10 min, this was held for 2 min, then decreased back to 40% B over 0.1 min, then held for 3 min. The retention time for 7α--hydroxy-4-cholesten-3-one was 8.2 min.

Data were acquired using SCIEX Analyst 1.6.2 and analyzed using SCIEX Multiquant 3.0.3 software. Sample values were calculated from calibration curves generated from the peak area ratio of the analyte to internal standard versus analyte concentration that was fit to a linear equation with 1/x weighting.

### FGF19 ELISA

Serum concentration of FGF-19 was determined using the Human FGF-19 Quantikine ELISA (R&D Systems, DF1900). Assay was completed according to manufacturer’s instructions with samples ran in duplicate.

#### Rodent Studies

### Postpartum rodent model

The University of Colorado-Anschutz Medical Campus approved animal procedures. Age-matched Sprague-Dawley female rats (Harlan, Indianapolis, IN) were housed and bred as described *(29)*. For tissue collection, rats were euthanized across groups CO2 asphyxiation and cardiac puncture. Whole livers were removed, washed 3x in 1x PBS, and tissues weighed. Left lobes were fixed in 10% neutral buffered formalin (Anatech ltd) and processed for FFPE and caudate lobes were flash frozen on liquid nitrogen for protein and RNA extraction.

### Immunohistochemistry

Immunohistochemical (IHC) detection was performed as described *(30)*. Briefly, tissues were deparaffinized, rehydrated, and heat-mediated antigen retrieval was performed with EDTA for 5 minutes at 125^°^C. Primary antibodies used were: Ki67 (Neomarkers RM-9106-s, 1:50) for 2 hours at RT and Adipophilin (LS-B2168/34250 Lifespan Biosciences, 1:400) for 1 hour at RT. Secondary antibody was anti-rabbit (Agilent Envision+ K4003, RTU), used for Ki67 at 1 hour at RT and for Adipophilin at 30 minutes at RT. DAB chromogen (Agilent, K346889-2) with hematoxylin counter stain (Agilent, S330130-2) was used to visualize positive stain. Stained sections were scanned using the Aperio AT2 slide scanner (Leica Biosystems). Number of Ki67+ hepatocytes were counted in 5-1×1mm areas. Adipophilin signal quantification was performed by Aperio ImageScope v12.1.0.5029 as described previously *(31)*. All analyses were done by investigators blinded to group.

### Quantitative real-time RT-PCR (qPCR)

RNA was isolated from flash frozen rat liver for cDNA synthesis and qPCR. One microgram total RNA was used for reverse transcriptase (RT)-mediated synthesis of cDNA using Superscript II RT (Invitrogen) and random hexamer primers for *Cyp7a* and Superscript IV (Invitrogen) for *FXR*. Quantitative PCR for rat *Cyp7a* and reference gene *GAPDH* was performed using FastStart Essential DNA Green Master (Roche) in an Applied Biosystems theromocycler with 45 cycles of 95^°^C for 20 seconds, 60^°^C for 40 seconds, 72^°^C for 20 seconds. Rat primer sequences were: *Cyp7a*, forward CTGTCATACCACAAAGTCTTATGTCA and reverse ATGCTTCTGTGTCCAAATGCC; *GAPDH* forward CGCTGGTGCTGAGTATGTCG and reverse CTGTGGTCATGAGCCCTTCC.

Quantitative PCR for rat *FXR* and reference gene *GAPDH* was performed using SsoAdvanced Unviersal SYBR Green Supermix (BioRad) in the ViiA 7 Real-Team PCR System (Thermo Fisher) with the following times: 95^?^C for 2 minutes, 40 cycles of 95^°^C for 15 seconds and 56^°^C for 60 seconds, then 95^°^C for 15 seconds, 61^°^C for 60 seconds, and 95^°^C for 15 seconds. Rat primer sequences were: *FXR*, forward AGGCCATGTTCCTTCGTTCA and reverse TTCAGCTCCCCGACACTTTT; *GAPDH*, forward ACCACAGTCCATGCCATCAC and reverse TCCACCACCCTGTTGCTGTA.

## Acknowledgements

We want to thank Mara Kalter and Claire Dorfman for participant outreach and scheduling, and to our study participants, whose generous gift of time and trust made this study possible. We are grateful to Dr. Gordon Mills for constructive review of the manuscript, and Weston Anderson for outstanding assistance with manuscript writing, editing, and preparation.

---

**Supplemental Table 1.**

|  | Full Liver Volume<br>Population<br>(N=47) | Liver Volume<br>measured at<br>pregnancy visits<br>only<br>(N=30) | Liver volume<br>measured at<br>pregnancy and<br>postpartum visits,<br>(N=17) |
| --- | --- | --- | --- |
|  | N (%) | N (%) | N (%) |
| **Age (years)** |  |  |  |
| mean (SD) | 30.2 (4.6) | 29.6 (4.7) | 31.2 (4.4) |
| [min-max] | [19.0-39.0] | [19.0-39.0] | [22.0-38.0] |
| **Preconception BMI (kg/mg<sup>2</sup>)** |  |  |  |
| mean (SD) | 25.9 (4.4) | 26.4 (4.2) | 25.2 (4.7) |
| [min-max] | [18.4-35.9] | [18.4-35.6] | [20.0-35.9] |
| Normal or Overweight (<30) | 40 (85.1) | 26 (86.7) | 14 (82.4) |
| Obese (≥30) | 7 (14.9) | 4 (13.3) | 3 (17.6) |
| **BMI at late pregnancy (kg/m<sup>2</sup>)** |  |  |  |
| mean (SD) | 29.5 (4.1) | 30.4 (3.7) | 28.0 (4.3) |
| [min-max] | [22.9-38.9] | [22.9-38.9] | [22.9-37.1] |
| Normal or Overweight (<30) | 27 (57.4) | 15 (50.0) | 12 (70.6) |
| Obese (≥30) | 20 (42.6) | 15 (50.0) | 5 (29.4) |
| **Parity** |  |  |  |
| 0 | 28 (59.6) | 21 (70.0) | 7 (41.2) |
| 1 | 10 (21.3) | 3 (10.0) | 7 (41.2) |
| >1 | 9 (19.1) | 6 (20.0) | 3 (17.6) |
| **Race** |  |  |  |
| White only | 40 (85.1) | 25 (83.3) | 15 (88.2) |
| Multiple races | 3 (6.4) | 1 (3.3) | 2 (11.8) |
| Unknown | 4 (8.5) | 4 (13.3) |  |
| **Ethnicity** |  |  |  |
| Hispanic | 6 (12.8) | 5 (16.7) | 1 (5.9) |
| Non-Hispanic | 41 (87.2) | 25 (83.3) | 16 (94.1) |
| **Gestational Age at Early Pregnancy (weeks)** |  |  |  |
| mean (SD) | 15.6 (0.8) | 15.4 (0.8) | 16.1 (0.8) |
| [min-max] | [12.9-17.6] | [12.9-16.6] | [14.7-17.6] |
| **Gestational Age at Late Pregnancy (weeks)** |  |  |  |
| mean (SD) | 34.3 (1.5) | 34.5 (1.5) | 33.8 (1.3) |
| [min-max] | [31.6-37.7] | [32.3-37.7] | [31.6-36.1] |
| **Gestational hypertension/pre-eclampsia** |  |  |  |
| Yes | 6 (12.8) | 2 (6.7) | 4 (23.5) |
| No | 41 (87.2) | 28 (93.3) | 13 (76.5) |
| **Intrahepatic cholestasis of pregnancy** |  |  |  |
| Yes | 2 (4.3) | 2 (6.7) | 0 |
| No | 45 (95.7) | 28 (93.3) | 17 (100.0) |
| **Newborn weight (kg)** |  |  |  |
| mean (SD) | 3.4 (0.5) | 3.4 (0.4) | 3.5 (0.7) |
| [min-max] | [2.0-4.5] | [2.6-4.5] | [2.0-4.5] |
| **Newborn length (cm)** |  |  |  |
| mean (SD) | 51.1 (2.8) | 51.3 (2.4) | 50.9 (3.4) |
| [min-max] | [44.0-56.0] | [47.0-56.0] | [44.0-56.0] |
| **Ponderal index (kg/m<sup>3</sup>)** |  |  |  |
| mean (SD) | 25.5 (2.7) | 25.2 (2.9) | 26.0 (2.5) |
| [min-max] | [20.4-32.1] | [20.4-32.1] | [21.9-29.6] |

**Supplemental Table 2.**

| Interoperator Variability in Liver Volume Measurement |  |  |  |  |
| --- | --- | --- | --- | --- |
| Case | Output | Operator #1 | Operator #2 | Difference between operator #1 and #2 |
| 100283 | Early Pregnancy (cm<sup>3</sup>) | 1142 | 1143 | 1 |
|  | Late Pregnancy (cm<sup>3</sup>) | 1376 | 1341 | -35 |
|  | Late-Early (cm<sup>3</sup>) | 234 | 198 | -36 |
|  | % Change | 20.5 | 17.3 | -3.2 |
| 100445 | Early Pregnancy (cm<sup>3</sup>) | 948 | 986 | 38 |
|  | Late Pregnancy (cm<sup>3</sup>) | 982 | 999 | 17 |
|  | Late-Early (cm<sup>3</sup>) | 34 | 13 | -21 |
|  | % Change | 3.6 | 1.3 | -2.2 |
| 100515 | Early Pregnancy (cm<sup>3</sup>) | 1203 | 1268 | 65 |
|  | Late Pregnancy (cm<sup>3</sup>) | 1727 | 1709 | -18 |
|  | Late-Early (cm<sup>3</sup>) | 524 | 441 | -83 |
|  | % Change | 43.6 | 34.8 | -8.7 |
| 100662 | Early Pregnancy (cm<sup>3</sup>) | 1015 | 1026 | 11 |
|  | Late Pregnancy (cm<sup>3</sup>) | 971 | 966 | -5 |
|  | Late-Early (cm<sup>3</sup>) | -44 | -60 | -16 |
|  | % Change | -4.33 | -5.89 | -1.55 |
| 101089 | Early Pregnancy (cm<sup>3</sup>) | 1300 | 1322 | 22 |
|  | Late Pregnancy (cm<sup>3</sup>) | 1835 | 1816 | -19 |
|  | Late-Early (cm<sup>3</sup>) | 535 | 494 | -41 |
|  | % Change | 41.2 | 37.3 | -3.8 |
|  |  | Average Difference in % Change |  | -3.9 |
|  |  | Standard Deviation in % Change |  | 2.8 |

**Supplemental Table 3.**
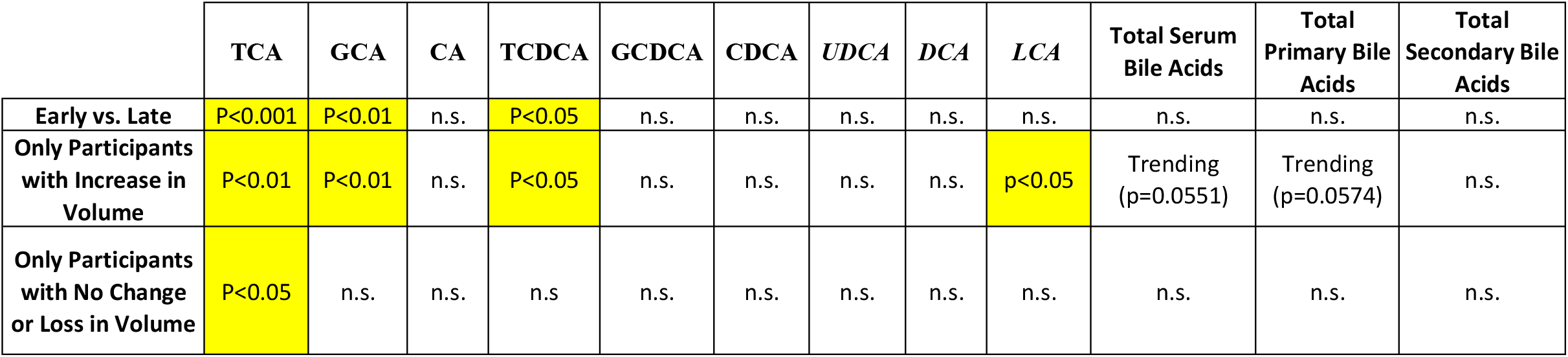

**SFig 1.**
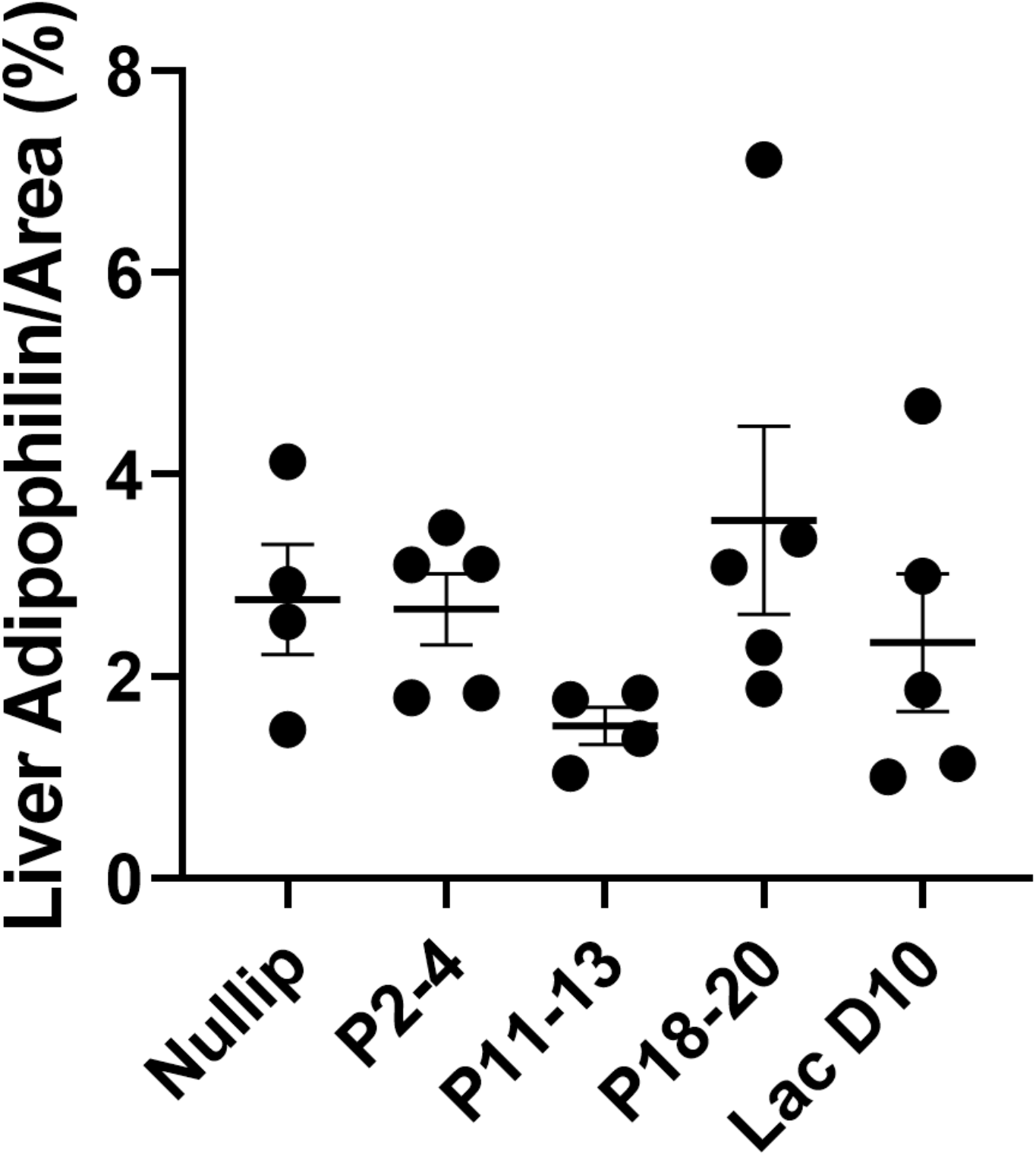
Quantification of adipophilin IHC staining in rat livers, n=4-5/group.

**SFig 2.**
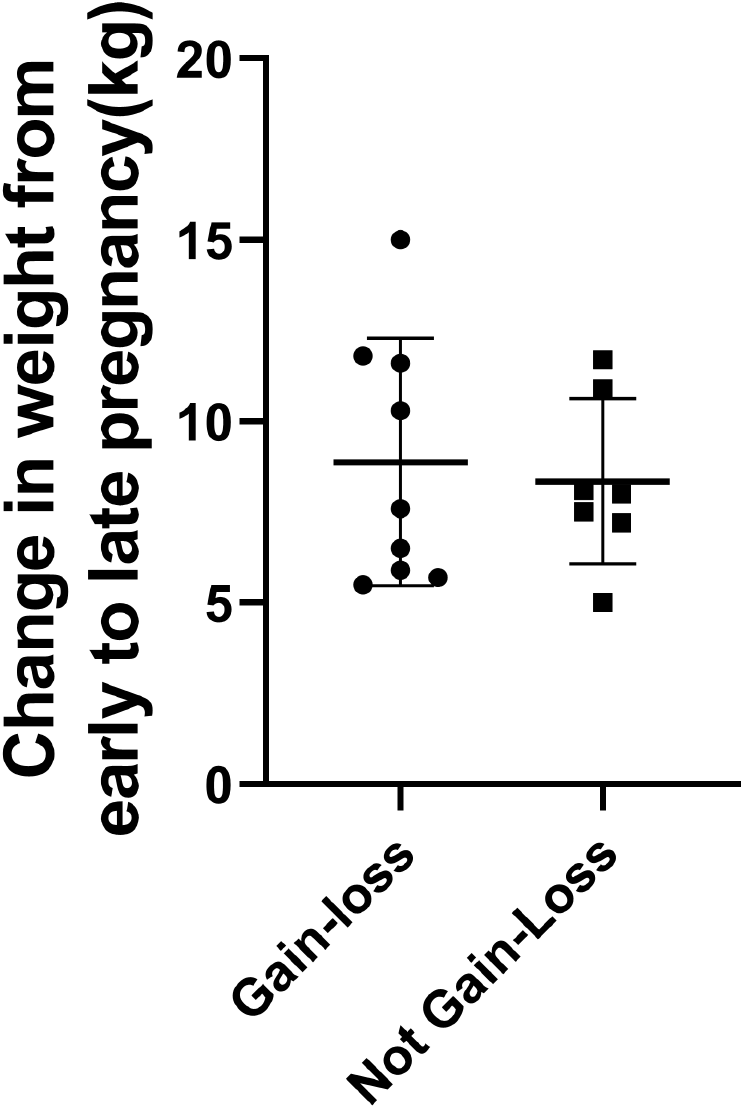
Change in weight from early to late pregnancy in participants who completed all 3 study visits, separated by whether their liver volume followed the gain-loss or not gain-loss pattern during pregnancy and post-wean.

**SFig 3.**
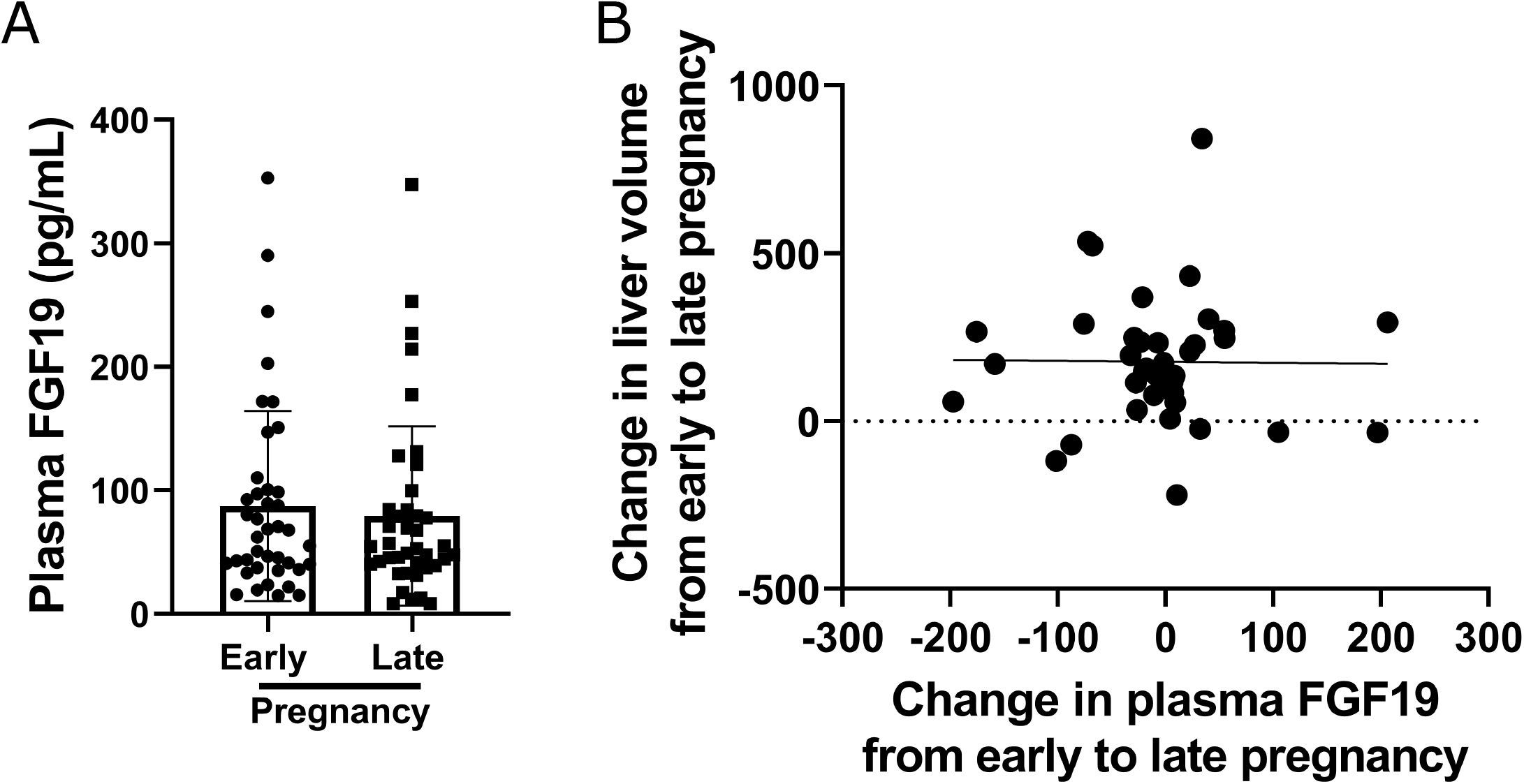
FGF19 Quantification. (A) Quantification of plasma FGF19 by ELISA at early and late pregnancy time point. (B) Pearson’s correlation of change in FGF19 and change in liver volume.

